# Compartmentalized SARS-CoV-2 replication in upper versus lower respiratory tract after intranasal inoculation or aerosol exposure

**DOI:** 10.1101/2023.09.11.557190

**Authors:** Robert J Fischer, Trenton Bushmaker, Brandi N. Williamson, Lizzette Pérez-Pérez, Friederike Feldmann, Jamie Lovaglio, Dana Scott, Greg Saturday, Heinz Feldmann, Vincent J. Munster, Emmie de Wit, Neeltje van Doremalen

## Abstract

Non-human primate models are essential for the development of vaccines and antivirals against infectious diseases. Rhesus macaques are a widely utilized infection model for severe acute respiratory syndrome coronavirus 2 (SARS-CoV-2). We compared cellular tropism and virus replication in rhesus macaques inoculated with SARS-CoV-2 via the intranasal route, or via exposure to aerosols. Intranasal inoculation results in replication in the upper respiratory tract and limited lower respiratory tract involvement, whereas exposure to aerosols results in infection throughout the respiratory tract. In comparison to multi-route inoculation, the intranasal and aerosol inoculation routes result in reduced SARS-CoV-2 replication in the respiratory tract.

---

The coronavirus disease 2019 (COVID-19) pandemic, caused by severe acute respiratory syndrome coronavirus 2 (SARS-CoV-2), has caused close to 7 million deaths as of the end of August 2023. The virus is transmitted primarily via aerosols and respiratory droplets [1]. Infected individuals can shed virus before exhibiting symptoms, with peak shedding detected at symptom onset [2]. Additionally, the virus has a half-life of over one hour as a suspended aerosol [3], which, unlike small droplets, can remain suspended for an extended period of time, suggesting that transmission can occur even after an infected person has left.

Animal models are instrumental in the investigation of pathogenesis caused by infectious agents, as well as in preclinical research on vaccines and antivirals [4]. However, in selecting an animal model, one must carefully consider the research question and weigh variables such as the inoculation route and resulting disease profile, to ensure that the studies are done in a consistent and stringent manner. We previously showed that the inoculation route affects SARS-CoV-2 disease progression in Syrian hamsters [5]. SARS-CoV-2 inoculation of non-human primate (NHP) models has been done via a multitude of different routes, including intranasal, intratracheal, and aerosol inoculation, with varying outcomes regarding disease severity, virus shedding, and tissue tropism [6–10]. In the current study, we directly compare SARS-CoV-2 tropism in rhesus macaques inoculated via the intranasal or aerosol route and compare outcomes to one of our previous NHP studies, in which we used a multi-route inoculation, combining the intranasal, intratracheal, oral, and ocular route [11].

## Methods

### Biosafety and ethics approval

Approval for studies involving NHPs was provided by the Animal Care and Use Committee at the Rocky Mountain Laboratories, Division of Intramural Research, National Institute of Allergy and Infectious Diseases, National Institutes of Health. Animal studies were carried out in an Association for Assessment and AAALAC International-accredited facility, following the basic principles and guidelines in the Guide for the Care and Use of Laboratory Animals, the Animal Welfare Act, US Department of Agriculture and the US Public Health Service Policy on Humane Care and Use of Laboratory Animals. Rhesus macaques were housed in individual primate cages enabling social interactions in a climate-controlled room with a fixed light/dark cycle (12-h/12-h). Commercial monkey chow, treats and fruit were provided by trained personnel. Water was available *ad libitum*. Environmental enrichment consisted of a variety of human interactions, manipulanda, treats, videos, and music. Animals were observed at least twice daily. The Institutional Biosafety Committee (IBC) approved work with infectious SARS-CoV-2 virus strains under BSL3 conditions at minimum. Virus inactivation of all samples was performed according to IBC-approved standard operating procedures for the removal of specimens from high containment areas.

### Cells and virus

SARS-CoV-2 strain SARS-CoV-2/human/USA/WA-001/2020 (MN985325.1 provided by the CDC) was obtained from a patient from Washington in 2020 and propagated in Vero E6 cells in DMEM supplemented with 2% fetal bovine serum (FBS), 1 mM L-glutamine, 50 U/mL penicillin and 50 μg/mL streptomycin. Vero E6 cells were provided by R. Baric, University of North Carolina, and mycoplasma testing was performed monthly. The virus stock was sequenced and analyzed using Bowtie2 version 2.2.9, and no single nucleotide polymorphisms (SNPs), compared to the patient sample sequence, were detected. Virus titrations were performed by end point titration in VeroE6 cells, which were inoculated with 10-fold serial dilutions of virus in 96-well plates. Cells were incubated at 37°C and 5% CO2. Cytopathic effect was read 5 days later.

### Experimental design

Four male rhesus macaques (aged 3 years, 4.2-5.2kg) were inoculated with SARS-CoV-2/human/USA/WA-001/2020 via the aerosol route with an average dose of 1.5 x 10^3^ TCID_50_/animal (all animals received between 9 x 10^2^ and 2.6 x 10^3^ TCID_50_, Supplementary Table 1). Aerosol exposure was performed using the AeroMP aerosol management platform (Biaera technologies, USA). Briefly, anesthetized macaques were exposed to a single exposure whilst contained in a head-only exposure unit. Aerosol droplet nuclei were generated by a 3-jet collison nebulizer (CH technologies, USA) and ranged from 1-5 µm in size. A sample of 6 liters of air per min was collected during the <10 min exposure on the 47 mm gelatin filter (Sartorius, Germany). Post exposure, the filters were dissolved in 10 mL of prewarmed (37°C) DMEM containing 10% FBS. Infectious virus was titrated as described above and the aerosol concentration was calculated. The inhaled inoculum was estimated using the respiratory minute volume rates of the animals, determined prior to the exposure, using the head-out Buxco plethysmography unit and FinePointe software (Data Sciences International). In the second cohort, four male rhesus macaques (aged 3 years, 3.6-4.8kg) were inoculated with the same strain of SARS-CoV-2 via the intranasal route with a dose of 8 x 10^5^ TCID_50_/animal, using 0.5 mL per naris (total volume of 1 mL). Hereafter, animals were observed and scored daily by the same person blinded to study group allocation using a standardized method [6]. Clinical examinations were performed on 0-, 1-, 3-, 5- and 7-days post inoculation (DPI). Nasal and oropharyngeal swabs were collected on all days that examinations were performed. Bronchoalveolar lavage (BAL) was performed on 3 and 5 DPI.

### RNA extraction and quantitative reverse-transcription polymerase chain reaction

Up to 30 mg of tissue was homogenized in RLT buffer and RNA was extracted using the RNeasy kit (Qiagen), whereas RNA was extracted from bronchoalveolar lavage fluid and swabs using the QiaAmp Viral RNA kit (Qiagen), both according to the manufacturer’s instructions. A viral genomic RNA-specific assay [12] was used for the detection of viral RNA.

### Histology

Collected tissues were fixed for a minimum of seven days in 10% neutral-buffered formalin, and subsequently embedded in paraffin, processed using a VIP-6 Tissue Tek tissue processor (Sakura Finetek), and embedded in Ultraffin paraffin polymer (Cancer Diagnostics). Samples were sectioned at 5 μm, stained with haematoxylin and eosin, and evaluated by a board-certified veterinary pathologist blinded to study groups.

### Statistical analyses

To investigate significant differences between the three study groups, two-way ANOVA with the Geisser-Greenhouse correction followed by Tukey’s multiple comparisons test or the Kruskal-Wallis test followed by Dunn’s multiple comparisons test was conducted using Graphpad Prism version 9.3.1.

### Data availability

All data can be accessed on https://figshare.com/articles/journal_contribution/_b_Compartmentalized_SARS-CoV-2_replication_in_upper_versus_lower_respiratory_tract_after_intranasal_inoculation_or_aerosol_exposure_b_/24036612

## Results and Discussion

An increase in clinical score was observed over time in all SARS-CoV-2-inoculated animals and was defined as mild disease (Figure 1A, euthanasia is mandated at a clinical score of ≥35). The clinical score was driven by respiratory signs and decreased appetite (Supplementary Table 2). No significant difference was observed between the clinical scores of the study groups at any time point, nor when the area under the curve for the clinical score was determined (Figure 1B). We then compared clinical scores to those obtained from NHP study previously performed in our laboratory [11], in which animals were inoculated via a combination of intranasal, intratracheal, oral, and ocular route and received a total dose of 2.6 x 10^6^ TCID_50_ per animal. Animals that received SARS-CoV-2 via the multi-route displayed higher clinical scores at 3, 4, 5, and 6 DPI than animals exposed to aerosols (Figure 1A), which was also reflected in the area under the curve analysis (Figure 1B). Nasal swabs were obtained on 1, 3, 5, and 7 DPI, and the presence of genomic viral RNA was investigated. In nasal swabs, the highest amount of viral RNA was detected at 1 DPI in animals inoculated via the intranasal and multi-route and at 3 DPI in animals exposed to aerosols. A significant difference was noted at 1 DPI between the multi-route and aerosol-inoculated animals (Figure 1C). All animals had evidence of virus replication in nasal swabs, and no significant difference in the total amount of virus shed was detected between the groups (Figure 1D). BAL was performed on 3 and 5 DPI. Genomic viral RNA levels were low in BAL fluid in animals inoculated via the intranasal or aerosol route, and significantly higher in animals inoculated via the multi-route (Figure 1E). Genomic viral RNA was detected in the BAL fluid of all animals inoculated via the aerosol route but only in one out of four animals inoculated via the intranasal route (Animal IN1, Figure 1F). Interestingly, animal IN1 also had a consistently higher clinical score throughout the study (Figure 1B). Genomic viral RNA detected in tissues isolated 7 DPI was low and was limited to the upper respiratory tract in animals inoculated via the intranasal route but detected in the lower respiratory tract of 3 out of 4 animals exposed to aerosols and 6 out of 6 animals exposed via the multi-route (Figure 1G). Thus, although no differences between groups were detected in virus replication in the upper respiratory tract, virus replication in the lower respiratory tract was only observed in animals exposed to aerosols compared to those inoculated via the intranasal route.

**Figure 1.**
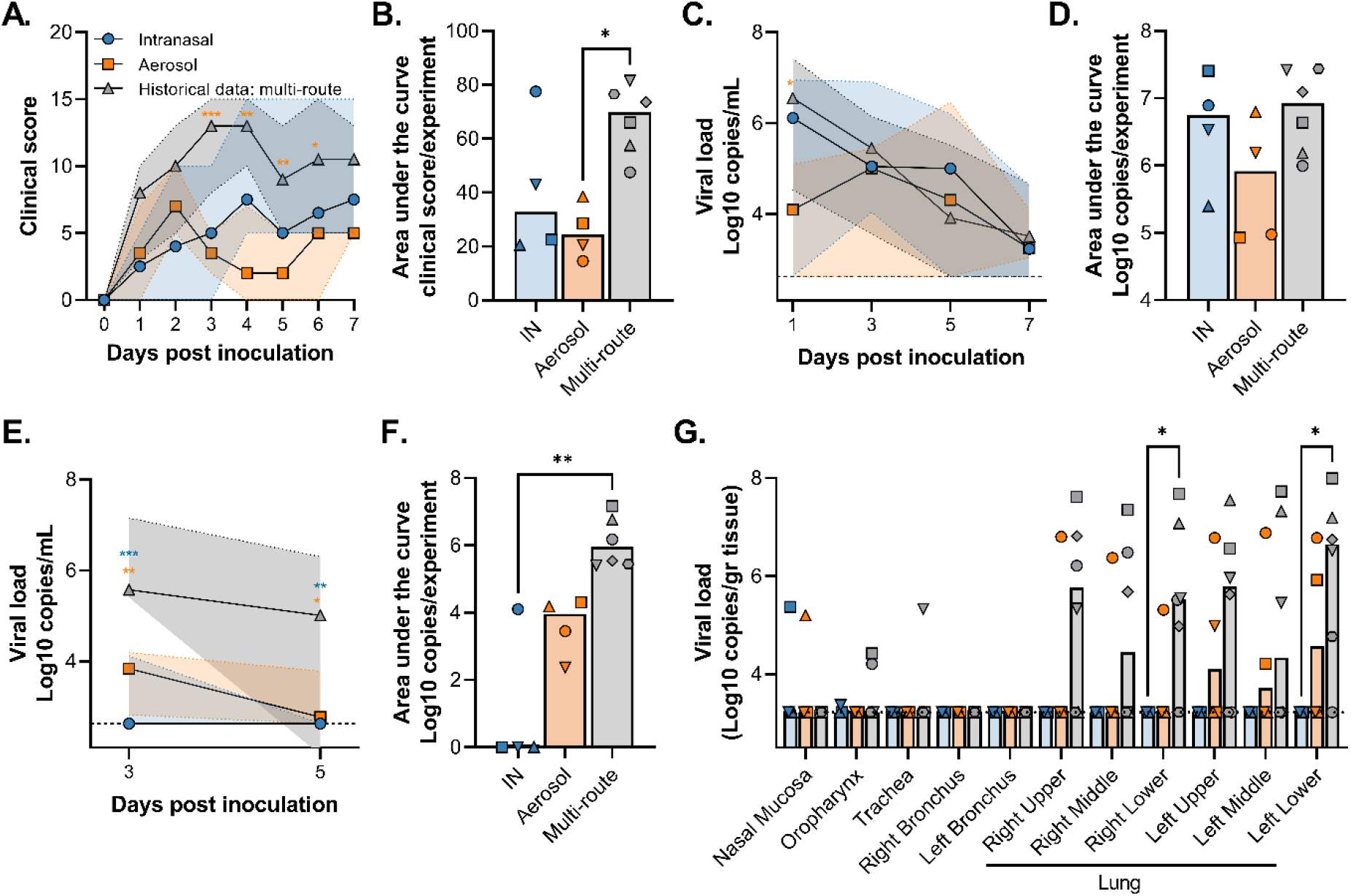
Clinical score and viral load in rhesus macaques after inoculation with SARS-CoV-2. Animals were inoculated via the intranasal route (blue), exposed to aerosols (orange), or inoculated via the intranasal, intratracheal, oral, and ocular route (grey, historical data from van Doremalen *et al*. [11]). A. Clinical signs were scored daily. Shown is median (symbols) with 95% confidence interval (shaded areas). B. Total clinical score for experiment. C, E. Genomic viral RNA detected in nasal swabs (C) and BAL fluid (E). Shown is median (symbols) with 95% confidence interval (shaded areas). Line = qualitative limit of detection. D, F. Total amount of genomic viral RNA shed per animal as measured in nasal swabs (D) and bronchoalveolar lavage fluid (F). Bar depicts median. I. Genomic viral RNA detected in tissue samples collected at 7 DPI. Bar depicts median. B, D, F. Circle: animal 1, Square: animal 2, upwards triangle: animal 3, downwards triangle: animal 4, diamond: animal 5, hexagon: animal 6. Statistical significance was determined via two-way ANOVA with the Geisser-Greenhouse correction followed by Tukey’s multiple comparisons test (A, C, E) or Kruskal-Wallis test followed by Dunn’s multiple comparisons test (B, D, F). Asterisks display group-specific differences, specified by color. *** = p-value < 0.001, ** = p-value <0.01, * = p-value <0.05.

Histological examination of lung tissue showed evidence of interstitial pneumonia in three out of four animals in both groups, defined as minimal lesions. The lesions in the lungs of two out of four animals exposed to aerosols included type 2 pneumocyte hyperplasia, which is typical for SARS-CoV-2 infection as described previously [6], whereas in accordance with the lack of SARS-CoV-2 replication in the LRT, type 2 pneumocyte hyperplasia was absent in the lung tissue of animals inoculated via the intranasal route (Table 1). When we used the multi-route to inoculate animals, type 2 pneumocyte hyperplasia was found in 3 out of 6 animals [11], suggesting a phenotype more like the aerosol exposed animals than the intranasally inoculated animals.

**Table 1.** Histological observations in NHPs inoculated via the intranasal or aerosol route. 0 = no lesions, 1 = minimal lesions (1-10%).

| Lung lobe | Observation | IN1 | IN2 | IN3 | IN4 | A1 | A2 | A3 | A4 |
| --- | --- | --- | --- | --- | --- | --- | --- | --- | --- |
| Right Upper | Pneumocyte hyperplasia | 0 | 0 | 0 | 0 | 1 | 0 | 0 | 0 |
|  | Interstitial pneumonia | 0 | 0 | 0 | 0 | 1 | 0 | 0 | 0 |
| Right Middle | Pneumocyte hyperplasia | 0 | 0 | 0 | 0 | 0 | 1 | 0 | 0 |
|  | Interstitial pneumonia | 0 | 0 | 0 | 0 | 0 | 1 | 0 | 0 |
| Right Lower | Pneumocyte hyperplasia | 0 | 0 | 0 | 0 | 0 | 0 | 0 | 0 |
|  | Interstitial pneumonia | 0 | 0 | 1 | 0 | 0 | 0 | 1 | 0 |
| Left Upper | Pneumocyte hyperplasia | 0 | 0 | 0 | 0 | 1 | 1 | 0 | 0 |
|  | Interstitial pneumonia | 0 | 1 | 1 | 1 | 1 | 1 | 1 | 0 |
| Left Middle | Pneumocyte hyperplasia | 0 | 0 | 0 | 0 | 0 | 1 | 0 | 0 |
|  | Interstitial pneumonia | 0 | 0 | 0 | 0 | 0 | 1 | 0 | 0 |
| Left Lower | Pneumocyte hyperplasia | 0 | 0 | 0 | 0 | 1 | 0 | 0 | 0 |
|  | Interstitial pneumonia | 0 | 0 | 0 | 1 | 1 | 0 | 0 | 0 |

Both infection routes resulted in mild and transient respiratory disease as defined by the clinical signs observed during the experiment. Importantly, no differences were observed in virus replication in the upper respiratory tract, but replication in the lower respiratory tract was more likely to occur in animals inoculated with aerosols as shown via viral RNA detection in BAL fluid and lung tissue. This is similar to previous work from our group, in which we show differences in disease progression in Syrian hamsters driven by the inoculation route: intranasal and aerosol inoculation was associated with more severe disease than fomite exposure [5].

Differences observed in this study are correlated with where the virus was deposited during inoculation; our data suggest that intranasal inoculation primarily deposited infectious virus in the upper respiratory tract, whereas aerosol and multi-route inoculation deposited virus in the entire respiratory tract, including the lower respiratory tract. This is in accordance with previous studies showing that inoculation route greatly affects deposition in the respiratory tract [13,14].

Although aerosol inoculation occurred with a much lower inoculum than intranasal inoculation (1.5 x 10^3^ vs. 8 x 10^5^ TCID_50_), no significant differences were found in upper respiratory tract replication. Likewise, in comparison with the multi-route inoculation, SARS-CoV-2 replication in the respiratory tract is reduced only in the lower respiratory tract, but not the upper respiratory tract. Analysis of 12 different influenza challenge studies in humans concluded that intranasal inoculation leads to 20 times lower infectivity compared to aerosol exposure [15], supporting this observation. It is possible that the bioavailability of aerosolized virus is much higher than that which is delivered in a mass volume. It should be noted that at 1 DPI, virus replication in the upper respiratory tract trends lower in the aerosol exposed group compared to the other two groups. This suggests that in animals exposed to aerosols, virus kinetics differ from the other two groups, but this did not affect the total amount of virus detected in nasal swabs. We have observed exposure-dependent SARS-CoV-2 shedding previously in Syrian hamsters [5].

In conclusion, we show that SARS-CoV-2 inoculation route utilized in rhesus macaque models affects the resulting virus tropism. Our findings support SARS-CoV-2 inoculation via the multi-route for the preclinical evaluation of vaccines and antivirals since this results in more virus replication in lung tissue and a robust phenotype. We recommend carefully considering these variables when selecting an animal model for subsequent studies.

## Author contributions

HF, VJM, EdW, and NvD designed the study; RJF, TB, BNW, LPP, FF, JL, GS, DS, VJM, EdW, and NvD acquired, analyzed, and interpreted the data; RJF and NvD wrote the manuscript. All authors have approved the submitted version.

## Funding

This work was supported by the Intramural Research Program of the National Institute of Allergy and Infectious Diseases (NIAID), National Institutes of Health (NIH).

## COI statement

The authors have no conflict of interest.

## Acknowledgements

We thank the animal caretakers of the Rocky Mountain Veterinary Branch, NIAID, NIH, and Myndi Holbrook, Laboratory of Virology, NIAID, NIH for their assistance during the study.

---

**Supplementary Table 1.** Infectious virus dose received per animal.

| Animal | Cumulative dose (TCID<sub>50</sub>) |
| --- | --- |
| IN1 | 8 x 10<sup>5</sup> |
| IN2 | 8 x 10<sup>5</sup> |
| IN3 | 8 x 10<sup>5</sup> |
| IN4 | 8 x 10<sup>5</sup> |
| A1 | 8.9 x 10<sup>2</sup> |
| A2 | 1.1 x 10<sup>3</sup> |
| A3 | 1.3 x 10<sup>3</sup> |
| A4 | 2.6 x 10<sup>3</sup> |
| M1 | 2.6 x 10<sup>6</sup> |
| M2 | 2.6 x 10<sup>6</sup> |
| M3 | 2.6 x 10<sup>6</sup> |
| M4 | 2.6 x 10<sup>6</sup> |
| M5 | 2.6 x 10<sup>6</sup> |
| M6 | 2.6 x 10<sup>6</sup> |

**Supplementary Table 2.** Clinical observations in animals inoculated with SARS-CoV-2. IN = intranasal; A = aerosol; M = multi-route.

| Animal | Day 0-3 | Day 4-7 |
| --- | --- | --- |
| IN1 | Increased abdominal respirations, Pale appearance, ruffled fur | Hunched posture, increased respirations, pale appearance, ruffled fur |
| IN2 | Irregular respirations | Irregular respirations |
| IN3 | Mildly decreased appetite | Irregular respirations |
| IN4 | Decreased appetite | Decreased appetite, irregular respirations |
| A1 | Clear nasal discharge, mildly irregular respirations, decreased appetite | Decreased appetite |
| A2 | Increased irregular respirations, clear nasal discharge, mildly ruffled fur | Increased irregular respirations, mildly ruffled fur |
| A3 | Deep abdominal respirations, decreased appetite | Lethargic, mildly irregular respirations, decreased appetite |
| A4 | Deep abdominal respirations, decreased appetite | mildly increased irregular respirations |
| M1 | Ruffled fur, decreased appetite, diarrhea, increased irregular breathing | Decreased appetite, diarrhea, increased irregular breathing |
| M2 | Ruffled fur, decreased appetite, increased irregular breathing | Ruffled fur, decreased appetite, increased irregular breathing |
| M3 | Ruffled fur, decreased appetite, increased abdominal breathing | Ruffled fur, decreased appetite, increased abdominal breathing |
| M4 | decreased appetite, pale appearance, hunched posture, deep abdominal & chest breathing | Hunched posture, decreased appetite, irregular abdominal breathing |
| M5 | Ruffled fur, decreased appetite, vomit, increased abdominal breathing | Decreased appetite, increased abdominal breathing, very agitated |
| M6 | Ruffled fur, decreased appetite, increased abdominal irregular breathing, pale appearance, very agitated | Ruffled fur, decreased appetite, increased abdominal irregular breathing, pale appearance, very agitated |

## Notes

https://figshare.com/articles/journal_contribution/_b_Compartmentalized_SARS-CoV-2_replication_in_upper_versus_lower_respiratory_tract_after_intranasal_inoculation_or_aerosol_exposure_b_/24036612

### Competing Interest Statement

The authors have declared no competing interest.

